# Axolotl tail regeneration emerges during a defined embryonic window

**DOI:** 10.64898/2026.05.21.726893

**Authors:** Anahi Binagui-Casas, Martin Nuamah Asare, Francisco Falcon, Valerie Wilson, Elly M. Tanaka, Wouter Masselink

## Abstract

How regenerative capacity originates during development remains poorly understood, even in vertebrates with exceptional adult regenerative ability. Using the axolotl, we identify a defined embryonic window between stages 30 and 34 during which the tail region transitions from a regeneration-incompetent to a regeneration-competent state. Amputations across staged embryos reveal that earlier embryos entirely fail to regenerate, whereas later embryos regenerate functional tails. Notably, tail stumps from nonregenerating embryos can recover the ability to regenerate when reamputated at later stages, demonstrating that early regenerative failure does not permanently impair regenerative capacity. This differs from the transient refractory period described in *Xenopus*, where regenerative competence is lost and reacquired around the end of tail outgrowth, and indicates that staged acquisition of regenerative competence is a broadly shared but mechanistically distinct feature of amphibian development. To determine whether this transition reflects changes in progenitor composition, we analysed the single-cell transcriptional landscapes of axolotl tail buds across this window. Tail bud progenitors, including neuromesodermal progenitors, persist through the transition, indicating that the onset of regenerative competence is unlikely to be explained by the loss of embryonic progenitors. Finally, using *Tbxt* (Brachyury) crispant axolotls with severe axial defects, we show that tail regeneration occurs effectively despite earlier abnormal embryonic tail development, with functional uncoupling of the mechanisms of tail development and regeneration. This framework provides new opportunities for identifying the drivers of regenerative competence and understand why this capacity is lost in other vertebrate species.

## Introduction

Some vertebrates have the ability to regenerate the tail in adulthood (Roy & Gatien, 2008), and amphibians have been key models for dissecting the cellular and molecular basis of this process (Chen et al., 2014; Wang et al., 2026). However, it remains unclear whether this capacity is present continuously from the earliest stages of development, or emerges later during embryogenesis, even in the axolotl (Reiss, 2022). This gap is notable given the extensive characterisation of larval and adult tail regeneration (Masselink et al., 2026; Romero et al., 2018; Tucker & Slack, 1995). Classical embryological studies have reported the suppression of tail regeneration at early tail bud stages in several amphibians, including axolotl (Münch, 1938; Schaxel, 1922; Svetlov, 1934; Vogt, 1931). In *Xenopus*, early suppression also occurs (Tucker & Slack, 1995), preceding the distinct refractory period at stages 45–47, representing a secondary, temporary loss of regenerative ability after competence has already been established (Aztekin et al., 2019; Beck et al., 2003; Tucker & Slack, 1995). Yet these early observations in axolotl were never revisited systematically, leaving the timing, reproducibility, and basis of the initial acquisition of regenerative competence unresolved.

A related question is the cellular basis of such transition into regenerative competency. Embryonic body axis elongation is driven by neuromesodermal progenitors (NMPs) in the tail bud (Wymeersch et al., 2021). In amniotes, these progenitors are depleted as development progresses. Because NMPs generate tissues that must be rebuilt during tail regeneration, and are relevant for cell-based regenerative therapies (Binagui-Casas et al., 2021), understanding whether regenerative competence is linked to their presence is an important unexplored question.

Here we report that axolotl tail buds amputated during a defined developmental window fail to regenerate, resulting in tail loss. Reanalysis of single-cell tail bud transcriptomes across regenerative and nonregenerative stages revealed that this transition is not explained by NMP loss. Moreover, neither aberrant embryonic tail morphogenesis nor complete tail absence prevents juvenile tail regeneration. These findings support the idea that tail development and regeneration are mechanistically distinct: the former driven by somitogenesis (and presumably by NMPs), and the latter by a progenitor population arising in juvenile animals without recapitulating development (Masselink et al., 2026). Together with other recent work on the thymus (Czarkwiani et al., 2025), these findings suggest that development and regeneration are separable modules, and that regenerative competence is an acquired state rather than a simple redeployment of developmental programmes in the tail.

## Results and Discussion

### 1. A defined developmental window of regenerative acquisition during axolotl embryogenesis

To test whether tail regenerative ability is present throughout axolotl embryogenesis, we amputated tail buds across staged embryos from early tailbud (st.26) through larval regenerating stages (st.43; (Ponomareva et al., 2015)) and assessed outcomes over six weeks (Figure 1A). Amputations were consistently performed at the cloacal axial level, a reproducible anatomical landmark ensuring the removal of *Sox2/Tbxt* coexpressing regions that are characteristic of NMPs (Taniguchi et al., 2017).

**Figure 1.**
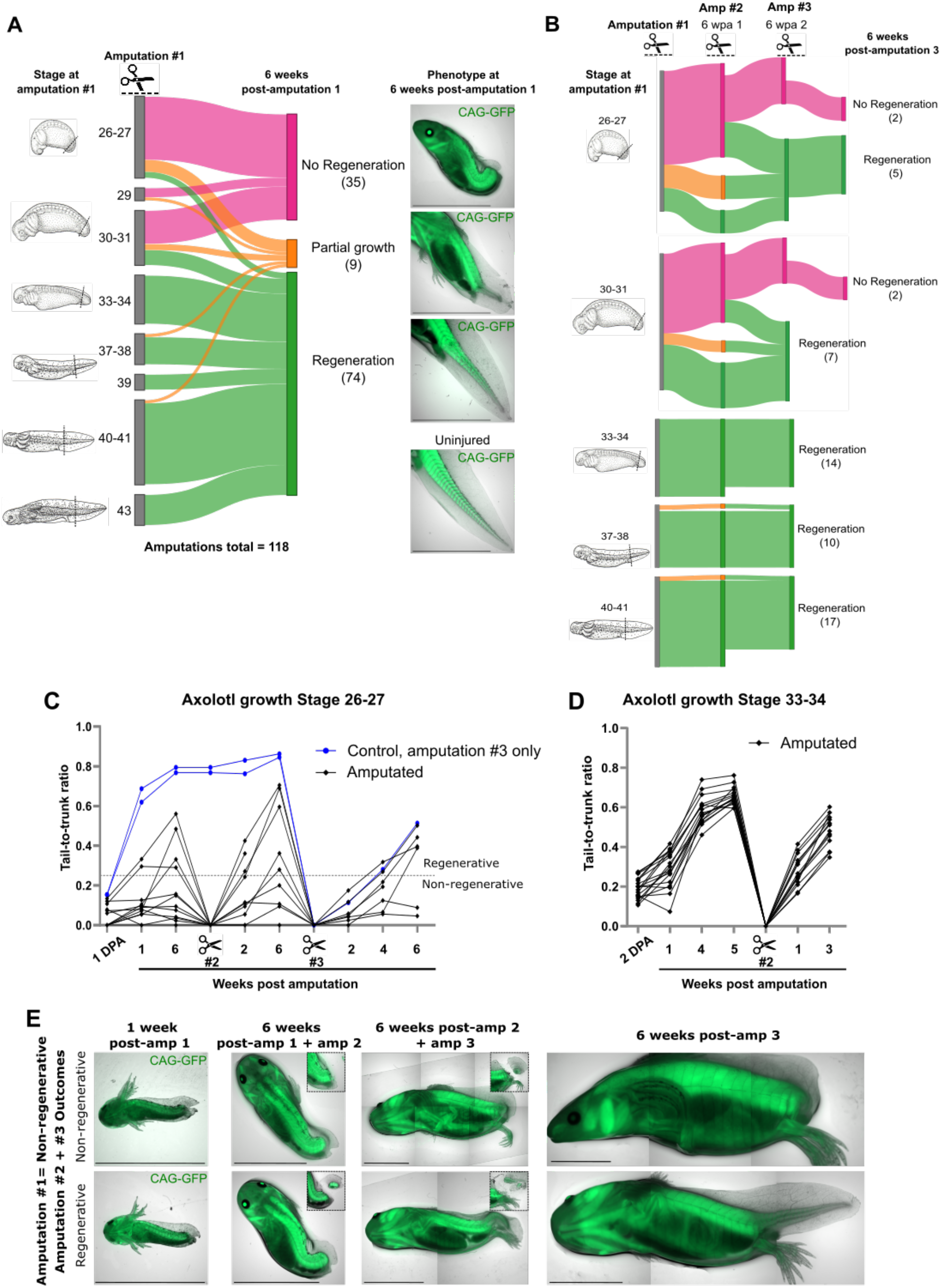
Axolotl embryos acquire tail regenerative competence within a defined developmental window. **A**. Sankey diagram showing regenerative outcomes following tail bud amputation across staged axolotl embryos (total n=118; CAG-GFP ubiquitous GFP reporter line). Amputations were performed at stages spanning early tail bud (st.26-27) through larval stages (st.43). The outcomes were scored at 6 weeks post-amputation as no regeneration (pink; n=35), partial growth (orange; n=9), or regeneration (green; n=74). Representative fluorescence images of each outcome phenotype are shown. A sharp transition in regenerative competence is observed between st.30-31 and st.33-34. **B**. Sankey diagram of secondary (6 wpa1) and tertiary (6 wpa2) amputations performed on primary stumps from animals across the same staged cohort, with final outcomes scored at 6 weeks post-tertiary amputation. Animals from nonregenerative embryonic stages (st.26-27: 5/7 regenerated; st.30-31: 7/9 regenerated) recover regenerative capacity upon repeated amputation challenge, whereas a small subset fail to regenerate persistently (st.26-27: n=2; st.30-31: n=2). Animals from st.33-34 onward show complete regeneration at all timepoints (st.33-34: n=14; st.37-38: n=10; st.40-41: n=17). Animals that did not survive to a full given amputation round were excluded from the plot. **C**. Tail-to-trunk ratio growth curves for animals amputated at st.26-27 (amputated, black; uninjured controls in Amp 1-2, blue), across three successive amputation rounds (Amp 1-3). Tail-to-trunk ratios remain consistently low in amputated relative to controls, confirming the absence of regenerative outgrowth at nonregenerative stages. **D**. Tail-to-trunk ratio growth curves for animals amputated at st.33-34, across two successive amputation rounds (Amp 1-2), showing robust and reproducible regenerative outgrowth following each amputation. **E**. Representative fluorescence images of CAG-GFP animals amputated at st.26-27 and their outcomes at secondary and tertiary amputations, with amputation planes in inset (top row, non-regenerating, bottom row regenerating). Scale bar: 1cm. Amp= amputation. wpa= weeks post-amputation.

Early-stage embryos (st.26-29) largely failed to regenerate (24/31 amputated; Figure 1A, E). These animals survived to adulthood, underwent wound closure, and developed normal anterior morphology, but showed no regenerative outgrowth and retained low tail-to-trunk ratios compared to uninjured controls (Figure 1C). In contrast, embryos amputated from st.33 onwards regenerated robustly, with functional tails observed in 67/69 animals by 6 weeks post-amputation (Figure 1A, D). Embryos amputated at intermediate stages (st.30-31) had mixed outcomes (11/18 non-regenerating, 5/11 regenerating and 2/11 partly regenerating), indicating a transition to regenerative competence between st.30-31 and st.33-34 (Figure 1A). This transition may reflect a graded acquisition of competence or minor technical/anatomical differences affecting amputation plane positioning.

We next addressed whether embryonic developmental history influences later regenerative capacity by performing secondary and tertiary amputations at 6-week intervals encompassing animals from both regenerative and nonregenerative primary cohorts (Figure 1B). Subsequent amputations were made distal to the cloaca for consistency and survival, although the low sample size and variable stump morphology in non-regenerators limited full standardisation. Strikingly, several animals from nonregenerative stages recovered regenerative capacity upon secondary and tertiary challenges: 5/9 st.26-27 primary non-regenerators regenerated after secondary amputation and persisted after tertiary amputation, while 3/7 st.30-31 animals regenerated after secondary amputation (Figure 1B). In contrast, animals amputated from st.33-34 onwards regenerated at all subsequent rounds (st.33-34: n=14; st.37-38: n=10; st.40-41: n=17; Figure 1B), indicating that once competence is established, it is robustly maintained. A small subset of early-stage animals failed to regenerate after repeated amputations (Figure 1B, E), suggesting that additional factors may modulate competence. Possible explanations include the malformation of key tissues such as the spinal cord (Schnapp et al., 2005; Sehm et al., 2009), or the permanent disruption of cloaca-adjacent signalling environments during primary amputation.

An uncoupling between embryonic development and adult regeneration was hinted at nearly a century ago (Svetlov, 1934) and is consistent with recent work which suggests axolotl tail regeneration does not recapitulate the somite-based development but instead relies on dermo-myo-sclerotome (DMS) progenitors (Masselink et al., 2026). Our data reveal a transition from regeneration incompetent to competent between st.30 and st.34, which is coherent with the findings of classical amphibian studies (Münch, 1938; Schaxel, 1922; Svetlov, 1934; Vogt, 1931). Repeated amputations further revealed that postembryonic regeneration is independent of complete tail formation during embryogenesis.

Developmental windows of regenerative competence are not unique to axolotl. In *Xenopus*, tail regeneration is transiently suppressed during a later refractory period (st.45-47) before competence is restored (Aztekin et al., 2019; Beck et al., 2003; Slack et al., 2004; Tucker & Slack, 1995). This period is associated with nutrient stress and yolk depletion (Williams et al., 2021). Interestingly, no equivalent incompetent stage has been identified in the axolotl limb bud, where repeated embryonic amputations instead produce permanently smaller limbs (Bryant et al., 2017), likely due to reduced innervation (Wells et al., 2021). Together, these findings suggest that axolotl tail regenerative incompetence reflects a local tissue state rather than a systemic constraint, and the staged acquisition of regenerative competence may be broadly conserved across amphibians.

### 2. Regenerative competence is acquired despite maintenance of embryonic progenitor identity

Having established that regenerative competence is acquired within a defined developmental window, we next asked what underlies this transition at the cellular level. Given the central role of NMPs in vertebrate tail morphogenesis, one possibility is that regenerative competence reflects a loss of NMPs or a shift in the progenitor state. To test this hypothesis, we reanalysed a published single-cell RNA-sequencing dataset of axolotl tail buds spanning the regenerative transition ((Masselink et al., 2026); see Methods).

Putative NMPs (*Sox2+/Tbxt+*), mesodermal derivatives (*Tbxt+/Sox2-*), and neural populations (*Sox2+/Tbxt-*) were detected at all stages (Figure 2A, B). Importantly, NMP-like populations were present at both regeneration-incompetent and regeneration-competent stages, with broadly comparable transcriptomic states based on well-known NMP markers (Figure 2B, C). Thus, acquisition of regenerative competence is not accompanied by NMP loss or a major change in progenitor identity, arguing against depletion of the embryonic axial progenitor pool as the primary driver of this transition.

**Figure 2.**
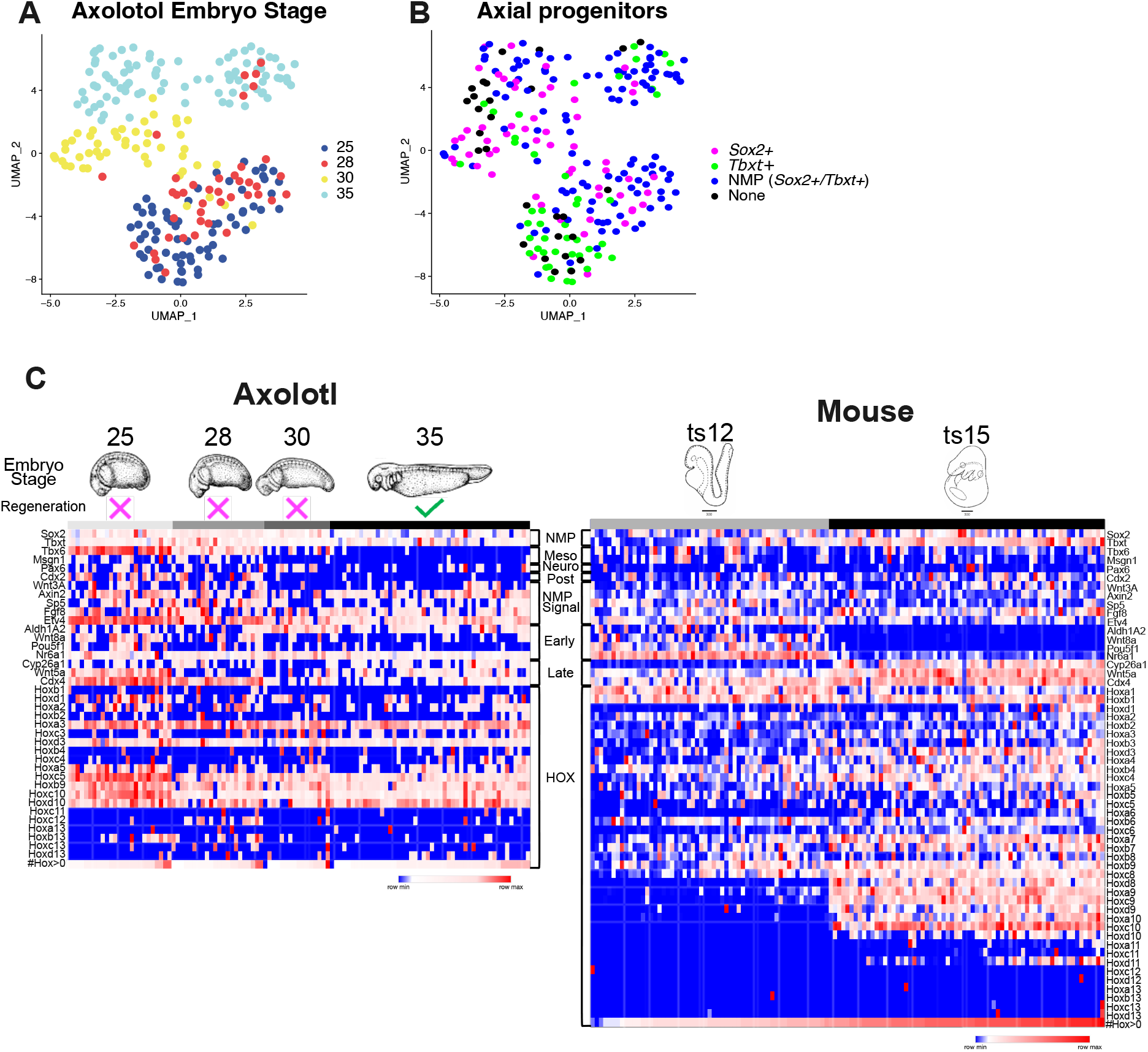
Single-cell RNA-seq analysis of axolotl tail buds across the regenerative transition. **A**. UMAP of axolotl tail bud single-cell RNA-seq data (Masselink et al., 2026), coloured by embryonic stage (st.25, st.28, st.30, st.35), spanning regeneration-incompetent and regeneration-competent timepoints. **B**. Same UMAP as in **A**, coloured by axial progenitor identity, based on *Sox2* and *Tbxt* coexpression: *Sox2*+ only (magenta), *Tbxt*+ only (green), NMP (*Sox2*+/*Tbxt*+ double-positive, blue), unassigned (black). NMP-like cells are present across all stages represented. **C**. Heatmap of NMP-associated and HOX gene expression across preselected NMPs (*Sox2* and *Tbxt*+ expression ≥1) in axolotl tail buds (st.25-35, left) and comparison with scRNA-seq of microdissected mouse tailbud NMPs (ts12-ts15, right) (Gouti et al., 2017), with embryo stage schematics and axolotl regenerative competency shown above each dataset. The following gene categories are annotated: NMP core genes (NMP), mesodermal (Meso), neural (Neuro), early and late progenitor markers (Early, Late), NMP signalling (NMP Signal), and HOX genes (HOX). Colour scale: row-normalised expression (blue = min, red = max expression). Data is ordered by stage and then by the total number of HOX genes with detectable expression.

Classical mouse NMP markers were also expressed in equivalent axolotl cells preselected for *Sox2/Tbxt* coexpression (Figure 2C). Comparison with a published mouse NMP single-cell dataset (Gouti et al., 2017) revealed broad transcriptional similarity with some exceptions. First, unlike mouse NMPs, which maintain posterior elongation programs throughout axis development, axolotl NMPs progressively downregulate tail elongation and mesoderm specification genes (Tbx6, Msgn1, Cdx2, Wnt3a) as regenerative competence emerges. Second, whereas mouse NMPs show progressive temporal HOX activation consistent with anterior-to-posterior collinearity across Theiler Stage (TS) 12-15, before NMP depletion by TS22 (Wymeersch et al., 2019), we found that axolotl NMPs do not show equivalent stage-dependent progressive HOX activation (st.25-35).

Instead, multiple HOX genes, including those in posterior paralogue group 13, were expressed across stages without clear temporal progression (Figure 2C; see row ‘#Hox>0’). These findings suggest that axolotl NMPs lack the late HOX maturation programme characteristic of mouse axial elongation. Whether this reflects a fundamentally different mode of NMP organisation in axolotl, whose axis continues elongating and segmenting well beyond hatching (Vincent et al., 2015), or limitations in cross-species staging equivalence remains unresolved. Nevertheless, these findings raise the possibility that salamander NMPs are regulated differently from their mammalian counterparts to sustain prolonged axial growth.

Collectively, our data show that the embryonic axolotl tail bud retains NMPs throughout the regenerative transition and that the acquisition of regenerative competence cannot be explained by loss or drastic identity changes in NMP-like progenitor states.

### 3. Tbxt is functionally dispensable for tail regeneration in chimeric F0 mutants

Our findings indicate that the acquisition of regenerative competence is not explained by changes in NMP identity. To test whether NMPs themselves are required for tail regeneration, we generated *Tbxt* F0 crispant axolotls using CRISPR/Cas9 mutagenesis. The axolotl genome encodes two *Tbxt* paralogues (on chromosomes 4q and 11q) that cluster with vertebrate *Tbxt* orthologues (Figure 3A). Based on synteny and sequence similarity, chromosome 4q likely harbours the canonical *Tbxt* and gRNAs were generated against a region in exon 2 (Flowers et al., 2014). *Tbxt* encodes a conserved T-box transcription factor that is required for posterior body axis elongation across vertebrates, is expressed in NMPs and is essential for their maintenance (Wymeersch et al., 2021), and its loss typically causes posterior truncation (Showell et al., 2004). We therefore assessed tail development and regeneration in F0 crispants and sibling controls.

**Figure 3.**
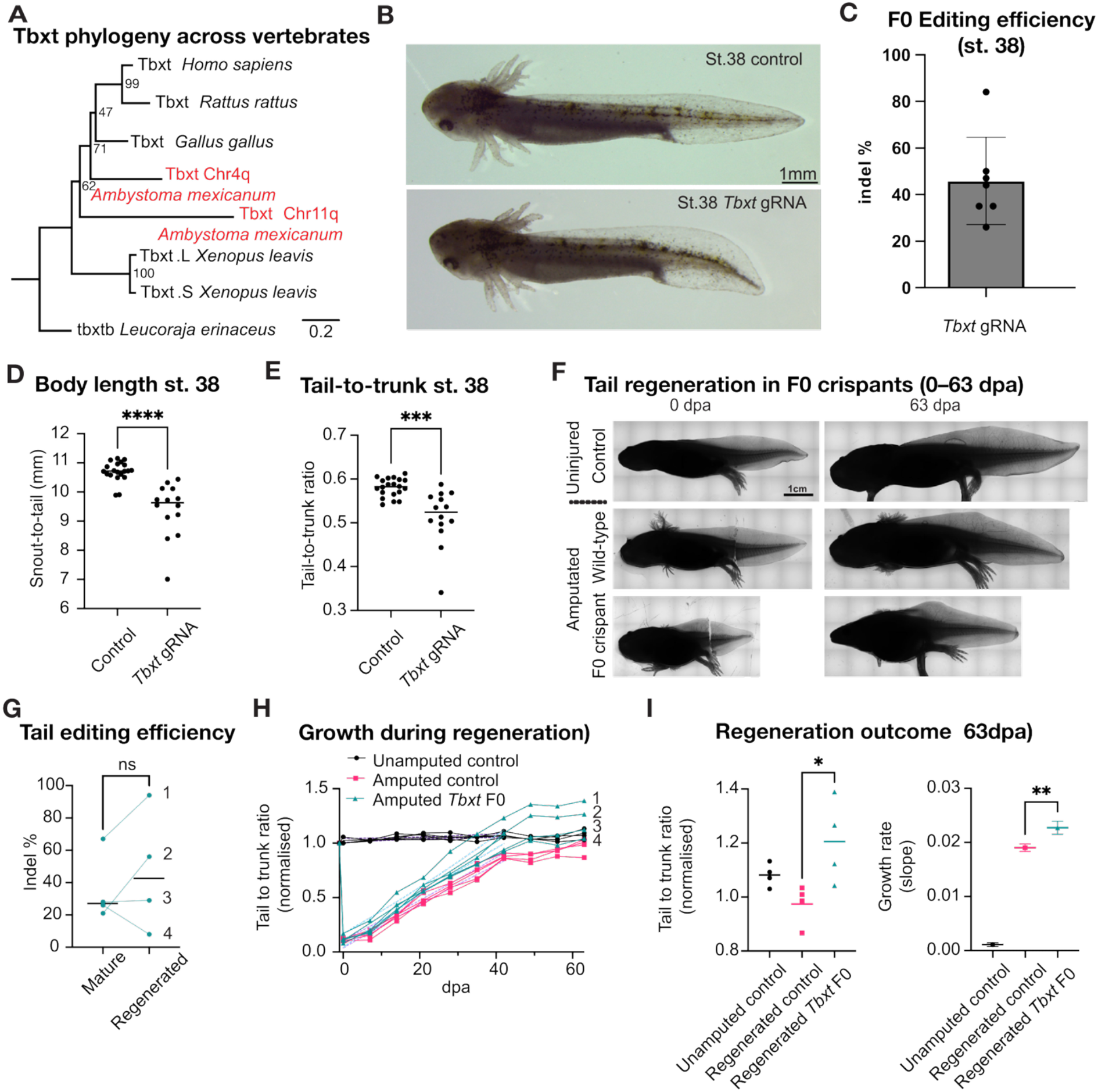
Tail regeneration proceeds in Tbxt F0 crispants despite severe defects in developmental tail outgrowth. **A**. Phylogenetic tree of Tbxt mRNA sequences across vertebrates, including two axolotl paralogous transcripts (from chromosomes 11q and 4q), *Xenopus laevis* L and S paralogues, and chicken, human, rat, and skate homologues. **B**. Representative brightfield images of st.38 control and *Tbxt* gRNA-injected crispant animals. Crispants display a reduction in tail length compared to controls. Scale bar: 1 mm. **C**. Indel percentage at the *Tbxt* target locus in F0 crispant animals, quantified by sequencing of tail tissue DNA. The mean indel rate was ~44%. **D**. Snout-to-tail-tip length at st.38 in control and *Tbxt* F0 crispant animals. Crispants are significantly shorter than controls. **E**. Tail-to-trunk ratio at st.38 in control and *Tbxt* F0 crispants. Crispants show a significant reduction in relative tail length, consistent with defects in posterior axis elongation. Variability in phenotype severity across crispants may correlate with variable indel rates shown in C. **F**. Representative brightfield images of uninjured control, amputated control (noncrispant), and amputated *Tbxt* F0 crispant animals at 0- and 63-days post-amputation (dpa). Both amputated controls and crispants regenerated tail structures by 63 dpa. Scale bar: 1 cm. **G**. Paired comparison of indel percentage at the *Tbxt* target locus between mature tail tissue and regenerated tail tissue in individual F0 crispant animals (n=4, numbered). No significant (ns) difference in editing efficiency was detected between timepoints. **H**. Normalised tail-to-trunk ratio over 63 dpa in unamputated controls (black), amputated controls (pink), and amputated *Tbxt* F0 crispants (blue). Crispants show robust regenerative outgrowth comparable to or exceeding amputated controls. **I**. Normalised tail-to-trunk ratio at 63 dpa (left) and regeneration growth rate (slope of tail-to-trunk ratio over time; right) across groups. Regenerated *Tbxt* F0 crispants have a significantly higher endpoint ratio and significantly higher growth rate than amputated controls, suggesting regeneration proceeds despite developmental tail defects. Statistical test: D, E: Unpaired t-test, F: Paired t-test, I: Ordinary one-way ANOVA. Lines and error bars represent mean ± SD. *p<0.05; **p<0.01; ***p<0.001; ****p<0.0001. dpa= days post amputation.

We were unable to recover homozygous *Tbxt* mutant axolotls, suggesting embryonic lethality. Mosaic F0 crispants that escaped lethality displayed pronounced tail elongation defects, consistent with previous reports (Flowers et al., 2014), including significantly reduced tail length at st. 38 (Figure 3 B-E) and in juvenile (7 cm Snout-to-tail in control siblings, Figure 3F) animals. Short-tail phenotypes here resemble chimeric mouse embryos containing *Tbxt* null and wildtype cells, where mutant NMPs accumulate at the tail bud preventing proper elongation (Guibentif et al., 2021; Wilson et al., 1995). Variability in phenotypic severity across individuals correlated with variable indel rates at the target locus (Figure 3C).

Strikingly, when *Tbxt* crispants were amputated at regeneration-competent stages, regeneration proceeded robustly despite impaired developmental growth. Regenerated tails recovered length and morphology comparable to those of the controls (Figure 3F), and when normalised to the snout-to cloaca length, the regenerated tails of crispants were longer than those of their wild-type siblings (Figure 3H, I). To exclude selective expansion of unedited cells, we compared editing efficiency between tissue adjacent to the amputation plane and regenerated tissue and found no significant difference (Figure 3G). This finding argues against strong selection for wild-type cells and suggests that *Tbxt* mutant cells do not disrupt differentiation during regeneration as they do during embryogenesis.

Together, our results demonstrate that tail regeneration can proceed independently of *Tbxt*-dependent embryonic axial growth, uncoupling a key developmental transcriptional regulator from the regenerative programme.

More broadly, our results demonstrate that regenerative competence in the axolotl tail emerges within a defined developmental window, is not associated with NMP loss, and is genetically uncoupled from *Tbxt*-dependent tail development. This is consistent with recent findings showing that adult axolotl tail regeneration depends on DMS progenitors contributing to all mesodermal lineages without recapitulating development (Masselink et al., 2026). However, regeneration is not solely driven by DMS progenitors. For example, spinal cord regeneration redeploys some developmental programmes, and is known to induce tail regeneration independently of DMS progenitors (Rodrigo Albors et al., 2015; Schnapp et al., 2005; Sehm et al., 2009). Thus, the degree of developmental programme reuse likely varies by tissue, while the acquisition of regenerative competence reflects a coordinated whole-tail transition that must coordinate tissue-specific programmes.

Similar distinctions have also been reported in axolotl thymus and zebrafish fin regeneration (Czarkwiani et al., 2025; Kang et al., 2016), supporting a broader model in which regeneration is not a simple reactivation of embryonic development, but relies on dedicated context-specific programmes.

The mechanisms underlying the acquisition of regenerative competence may be cell-autonomous or non-cell-autonomous. Epigenetic remodelling represents a plausible cell autonomous mechanism, as regeneration-associated loci in zebrafish are maintained in poised chromatin states that are activated upon injury (Stewart et al., 2009), with similar mechanisms proposed in amphibians (Katsuyama & Paro, 2011). Alternatively, and not mutually exclusive, injury-responsive cell populations such as DMS-like progenitors may not yet have acquired regenerative competence at early stages. Notably, these cells were not captured in the developmental scRNA-seq dataset analysed here. Non-cell-autonomous mechanisms, like immune signalling, may also contribute. For example, immune suppression during the refractory period in *Xenopus* partially restores regenerative competence (Fukazawa et al., 2009) whereas, the myeloid lineage itself is required for regeneration at the regenerative stages in *Xenopus* (Aztekin et al., 2020).

From an evolutionary perspective, these findings raise the possibility that regenerative programmes may function as distinct modules that can be differentially maintained, modified, or lost across species. Such modularity could help explain variation in regenerative capacity and implies that restoring complex structures may require activation of dedicated regenerative programmes rather than recapitulation of embryonic development. Comparative studies across diverse regenerative species may therefore reveal alternative cellular and molecular strategies for tissue restoration with translational potential, and the faithful reconstruction of complex structures in mammals may ultimately require the activation of such dedicated regenerative programmes rather than attempts to recapitulate embryonic development. Whether axolotl embryonic regeneration shares molecular and cellular features with juvenile and adult regeneration programmes remains an important open question, and one that is now tractable given the staging framework established in this study.

## Materials and Methods

### Animal Husbandry and Surgery

Axolotls (*Ambystoma mexicanum*) were bred in the Institute of Molecular Pathology facilities (Vienna, Austria.) All animal handling and surgical procedures were carried out in accordance with local ethics committee guidelines. Animal experiments were performed as approved by the Magistrate of Vienna. Animals under regulated procedures were anaesthetised with 0.03% benzocaine (Sigma-Aldrich) before amputation and surgery, and treated with the analgesic Butorphanol (Butomidor 0.5 mg/L, Vetviva Richter GmbH) for up to 3 days post-surgery. Embryonic amputations were performed under 0.03% benzocaine. Unless otherwise stated, we used “white” axolotls, which refers to a non-transgenic strain of axolotl (d/d) that has leucistic skin due to a melanocyte migration defect during embryonic development and carries an Edn3 mutation (Woodcock et al., 2017). CAG-GFP refers to the previously generated ubiquitous GFP reporter (Sobkow et al., 2006) with particularly high levels of GFP expression in the muscle.

Summary of embryonic experimental amputations:

**Table 1.** Animals underwent up to three sequential tail amputations. Amputations 2 and 3 were performed 6 weeks after the previous one. Stage refers to the NF stage at the time of the first amputation. Numbers decrease across successive amputations because of husbandry losses between rounds, unrelated to the amputation procedure itself; animals that did not survive to a full given amputation round are excluded from that column. n = number of individual animals; exp = number of independent experiments.

| Stage at 1st amputation | 1st amputation |  | 2nd amputation |  | 3rd amputation |  |
| --- | --- | --- | --- | --- | --- | --- |
|  | n | exp | n | exp | n | exp |
| 26–27 | 27 | 2 | 12 | 2 | 7 | 2 |
| 29 | 4 | 1 | NA | — | NA | — |
| 30–31 | 18 | 2 | 18 | 2 | 2 | 1 |
| 33–34 | 16 | 1 | 14 | 1 | NA | — |
| 37–38 | 10 | 1 | 10 | 1 | NA | — |
| 39 | 5 | 1 | NA | — | NA | — |
| 40–41 | 28 | 3 | 17 | 2 | NA | — |
| 43 | 10 | 1 | NA | — | NA | — |
| **Total animals** | **118** |  | **71** |  | **9** |  |

### Single cell RNA-seq analyses

The single-cell RNA-seq data was processed as described in Masselink et al. (2026). GEO accession number will be made available upon publication. We used Seurat (v5.3.1) for visualisation and differential gene expression analysis using the “FindMarkers” function. The low cell numbers in this dataset reflect the manual cell collection method used in the original study, where tail bud cells were individually picked for plate-based single-cell RNA-seq. While this limits the ability to identify rare cell types, it provides high read depth per cell (60,000 reads and a median of 6,731 genes per cell).

For the data shown in Figure 2C, axolotl cells were preselected for both *Sox2* and *Tbxt* expression, with at least 1 count of each gene, yielding 111 cells across stages 25, 28, 30, and 35. The mouse cells used for comparison were microdissected NMP cells across Theiler stages 12-15 from Gouti et al., 2017. Heatmap visualisation was performed using Morpheus, https://software.broadinstitute.org/morpheus, with row-normalised expression (z-score). Cells were ordered by embryonic stage and within stage by the total number of HOX genes with detectable expression (>0 counts).

### Phylogenetic analysis

*Tbxt* mRNA sequences for *Homo sapiens, Xenopus laevis, Gallus gallus*, and *Leucoraja erinacea* were obtained from the latest assemblies available in the UCSC genome browser (https://genome.ucsc.edu), and the *Ambystoma mexicanum Tbxt* sequences were obtained from the currently available annotation (AmexG_v6.0 (Schloissnig et al., 2021)). The sequences were aligned using CLUSTAL Omerga (v1.2.4) with default parameters. The multiple sequence alignment was manually processed by removing regions with large gaps and low sequence identity to other sequences, and the resulting alignment was used to construct a phylogenetic tree with IQ-TREE2 (using parameters: -m TEST -B 1000).

### Generation of *Tbxt* mutants

*Tbxt* mutants were generated according to Flowers et al. (2014) with modifications. Two primers containing a T7 promoter and *Tbxt* gRNA sequence (bolded) 5’-GAAATTAATACGACTCACTATAGG**AGGAATGCCTTCCGGTTT**GTTTTAGAGCTAGAAAT AGC-3’, and the ‘big reverse’ primer containing the sgRNA tracr sequence 5’-AAAAGCACCGACTCGGTGCCACTTTTTCAAGTTGATAACGGACTAGCCTTATTTTAACTT GCTATTTCTAGCTCTAAAAC-3’, were annealed, end-filled and purified using paramagnetic beads. We transcribed the sgRNAs with a MEGAscript T7 kit (Life Technologies). The template was digested using TURBO DNase (Thermo Fisher Scientific) and purified by lithium chloride precipitation. RNPs were generated by mixing sgRNA with in-house produced CAS9 protein (Fei et al., 2018). Freshly laid d/d eggs were injected with the RNP mixture and subsequently left to develop to the desired stage.

### Genotyping

Genotyping samples were lysed using 100 uL of 50 mM NaOH and incubated at 95°C for 15 minutes, neutralised using Tris, and centrifuged. The supernatant was moved to a clean tube, taking care not to disturb the debit. 1 µL of this stock was used for PCR genotyping. The sgRNA target site was amplified using flanking primers Tbxt_3UTR_Fwd (5’-AGCGTGGGTAGTTTTGATGG-3’) and Tbxt_3UTR_Rev (5’-CCGTGATGAGAAACGAGGTT-3’). PCR products for both wild-type and gRNA injected eggs were purified using paramagnetic beads and Sanger sequenced using the primer Tbxt_3UTR_Seq (5’-TCACTTTTTAAAGGGAGCACA-3’). Indel percentages were determined using Inference of CRISPR Edits (ICE) analysis (EditCo https://ice.editco.bio/#)

### Microscopy

Fluorescent and brightfield (black and white) imaging was performed using an Axio zoom.V16 (Zeiss, Jena) equipped with an Orca-Flash4.0 Camera (Hamamatsu Photonics, Hamamatsu) and an X-Cite Xylis LED illuminator (Excelitas Technologies, Waltham). Colour images were obtained using an SZx10 stereo-microscope (Olympus, Tokyo) equipped with a KL1600 LED swan-neck light (Olympus, Tokyo) and an AxioCam Erc 5s colour camera (Zeiss, Jena).

### Image and statistical analysis

Image analysis and measurements were performed in Fiji (Schindelin et al., 2012). Data flows were visualized using SankeyMATIC (www.sankeymatic.com). Statistical analysis and data visualisation were performed using Prism10.1.0 for Mac (GraphPad Software, Boston). Comparisons of control and *Tbxt* gRNA-injected larvae for both snout-to-tail tip and tail-to-trunk ratio was performed using a two-tailed unpaired t-test. Comparisons of indel% in mature and regenerated samples was performed using a two-tailed paired t-test. The normalised tail-to-trunk ratio was calculated by dividing the length of the tail (cloaca to tail tip) by that of the trunk (snout to cloaca), and normalised to the ratio calculated on −1 dpa (i.e., the day before amputation). Comparisons of the normalised tail to trunk ratio was performed using an ordinary one-way Anova followed by Tukey’s multiple comparisons test. The slope for each group was generated by performing a straight-line linear fit. Slope values were compared using an ordinary one-way ANOVA followed by Šídák’s multiple comparisons test, with error bars representing SD. ns: not significant, *: P ≤ 0.05, **: P≤ 0.01, ***: P≤ 0.001, ****: P≤0.0001. No statistical methods were used to predetermine sample size; all available animals from each independent experiment were included. n refers to the number of individual animals throughout, unless otherwise stated. ‘Independent experiments’ refers to clutches of embryos from separate parental crosses

## Author contributions

**Conceptualization:** A.B-C, V.W, W.M

**Methodology:** A.B-C, W.M

**Software:** F.F

**Validation:** A.B-C, W.M

**Formal analysis:** F.F, V.W, W.M

**Investigation:** A.B-C, M.N.A, W.M

**Resources:** W.M, E.M.T

**Data curation:** A.B-C, M.N.A, F.F, W.M

**Writing – original draft:** A.B-C

**Writing – review & editing:** A.B-C, W.M, V.W, F.F, M.N.A,

**Visualization:** A.B-C, W.M, V.W, F.F

**Supervision:** V.W, E.M.T

**Project administration:** A.B-C, W.M

**Funding acquisition:** A.B-C, W.M, E.M.T

## Acknowledgements

We would like to acknowledge the valuable services provided by the animal facility, Molecular Biology Services, and BioOptics facilities at IMP and IMBA. We would like to thank Yuka Taniguchi-Sugiura for help with the generation of *Tbxt* crispant axolotls. We would also like to thank Aida Rodrigo Albors and Sally Lowell for critically reading the manuscript and providing comments.

## Data availability

Data availability: All relevant data can be found within the article.

## References

Aztekin, C., Hiscock, T. W., Butler, R., De Jesus Andino, F., Robert, J., Gurdon, J. B., & Jullien, J. (2020). The myeloid lineage is required for the emergence of a regeneration-permissive environment following Xenopus tail amputation. Development, 147(3). 10.1242/dev.185496

Aztekin, C., Hiscock, T. W., Marioni, J. C., Gurdon, J. B., Simons, B. D., & Jullien, J. (2019). Identification of a regeneration-organizing cell in the Xenopus tail. Science, 364(6441), 653–658. 10.1126/science.aav9996

Beck, C. W., Christen, B., & Slack, J. M. (2003). Molecular pathways needed for regeneration of spinal cord and muscle in a vertebrate. Dev Cell, 5(3), 429–439. 10.1016/s1534-5807(03)00233-8

Binagui-Casas, A., Dias, A., Guillot, C., Metzis, V., & Saunders, D. (2021). Building consensus in neuromesodermal research: Current advances and future biomedical perspectives. Curr Opin Cell Biol, 73, 133–140. 10.1016/j.ceb.2021.08.003

Bryant, D. M., Sousounis, K., Farkas, J. E., Bryant, S., Thao, N., Guzikowski, A. R., Monaghan, J. R., Levin, M., & Whited, J. L. (2017). Repeated removal of developing limb buds permanently reduces appendage size in the highly-regenerative axolotl. Dev Biol, 424(1), 1–9. 10.1016/j.ydbio.2017.02.013

Chen, Y., Love, N. R., & Amaya, E. (2014). Tadpole tail regeneration in Xenopus. Biochem Soc Trans, 42(3), 617–623. 10.1042/BST20140061

Czarkwiani, A., Lobo, M., Castro, L. A. B., Petzold, A., Rost, F., Maehr, R., & Yun, M. H. (2025). Molecular basis for de novo thymus regeneration in a vertebrate, the axolotl. Sci Immunol, 10(114), eadw9903. 10.1126/sciimmunol.adw9903

Fei, J. F., Lou, W. P., Knapp, D., Murawala, P., Gerber, T., Taniguchi, Y., Nowoshilow, S., Khattak, S., & Tanaka, E. M. (2018). Application and optimization of CRISPR-Cas9-mediated genome engineering in axolotl (Ambystoma mexicanum). Nat Protoc, 13(12), 2908–2943. 10.1038/s41596-018-0071-0

Flowers, G. P., Timberlake, A. T., McLean, K. C., Monaghan, J. R., & Crews, C. M. (2014). Highly efficient targeted mutagenesis in axolotl using Cas9 RNA-guided nuclease. Development, 141(10), 2165–2171. 10.1242/dev.105072

Fukazawa, T., Naora, Y., Kunieda, T., & Kubo, T. (2009). Suppression of the immune response potentiates tadpole tail regeneration during the refractory period. Development, 136(14), 2323–2327. 10.1242/dev.033985

Gouti, M., Delile, J., Stamataki, D., Wymeersch, F. J., Huang, Y., Kleinjung, J., Wilson, V., & Briscoe, J. (2017). A Gene Regulatory Network Balances Neural and Mesoderm Specification during Vertebrate Trunk Development. Dev Cell, 41(3), 243–261 e247. 10.1016/j.devcel.2017.04.002

Guibentif, C., Griffiths, J. A., Imaz-Rosshandler, I., Ghazanfar, S., Nichols, J., Wilson, V., Gottgens, B., & Marioni, J. C. (2021). Diverse Routes toward Early Somites in the Mouse Embryo. Dev Cell, 56(1), 141–153 e146. 10.1016/j.devcel.2020.11.013

Kang, J., Hu, J., Karra, R., Dickson, A. L., Tornini, V. A., Nachtrab, G., Gemberling, M., Goldman, J. A., Black, B. L., & Poss, K. D. (2016). Modulation of tissue repair by regeneration enhancer elements. Nature, 532(7598), 201–206. 10.1038/nature17644

Katsuyama, T., & Paro, R. (2011). Epigenetic reprogramming during tissue regeneration. FEBS Lett, 585(11), 1617–1624. 10.1016/j.febslet.2011.05.010

Masselink, W., Gerber, T., Singh Jamwal, V., Falcon, F., Deshayes, T., Grindle, R., Seaman, R. P., Röcklinger, M., Papadopoulos, S.-C., Deneke, G., Adhikary, A., Andriotis, O. G., Pende, M., Taniguchi-Sugiura, Y., Lin, T.-Y., Kurth, T., Wang, J., Arendt, D., Fei, J.-F., … Murawala, P. (2026). Buckling instability underlies vertebral segmentation during axolotl tail regeneration. bioRxiv, 2024.2001.2031.577464. 10.1101/2024.01.31.577464

Münch, H. (1938). Über Regeneration in der Frühentwicklung: Defektoperationen im Gebiet der frühembryonalen Schwanzanlage bei Amphibien. Wilhelm Roux’Archiv für Entwicklungsmechanik der Organismen, 137(5), 597–635.

Ponomareva, L. V., Athippozhy, A., Thorson, J. S., & Voss, S. R. (2015). Using Ambystoma mexicanum (Mexican axolotl) embryos, chemical genetics, and microarray analysis to identify signaling pathways associated with tissue regeneration. Comp Biochem Physiol C Toxicol Pharmacol, 178, 128–135. 10.1016/j.cbpc.2015.06.004

Reiss, C. (2022). Cut and Paste: The Mexican Axolotl, Experimental Practices and the Long History of Regeneration Research in Amphibians, 1864-Present. Front Cell Dev Biol, 10, 786533. 10.3389/fcell.2022.786533

Rodrigo Albors, A., Tazaki, A., Rost, F., Nowoshilow, S., Chara, O., & Tanaka, E. M. (2015). Planar cell polarity-mediated induction of neural stem cell expansion during axolotl spinal cord regeneration. Elife, 4, e10230. 10.7554/eLife.10230

Romero, M. M. G., McCathie, G., Jankun, P., & Roehl, H. H. (2018). Damage-induced reactive oxygen species enable zebrafish tail regeneration by repositioning of Hedgehog expressing cells. Nat Commun, 9(1), 4010. 10.1038/s41467-018-06460-2

Roy, S., & Gatien, S. (2008). Regeneration in axolotls: a model to aim for! Exp Gerontol, 43(11), 968–973. 10.1016/j.exger.2008.09.003

Schaxel, J. (1922). Über die Natur der Formvorgänge in der tierischen Entwicklung. Archiv für Entwicklungsmechanik der Organismen, 50(3), 498–525.

Schindelin, J., Arganda-Carreras, I., Frise, E., Kaynig, V., Longair, M., Pietzsch, T., Preibisch, S., Rueden, C., Saalfeld, S., Schmid, B., Tinevez, J. Y., White, D. J., Hartenstein, V., Eliceiri, K., Tomancak, P., & Cardona, A. (2012). Fiji: an open-source platform for biological-image analysis. Nat Methods, 9(7), 676–682. 10.1038/nmeth.2019

Schloissnig, S., Kawaguchi, A., Nowoshilow, S., Falcon, F., Otsuki, L., Tardivo, P., Timoshevskaya, N., Keinath, M. C., Smith, J. J., Voss, S. R., & Tanaka, E. M. (2021). The giant axolotl genome uncovers the evolution, scaling, and transcriptional control of complex gene loci. Proc Natl Acad Sci U S A, 118(15). 10.1073/pnas.2017176118

Schnapp, E., Kragl, M., Rubin, L., & Tanaka, E. M. (2005). Hedgehog signaling controls dorsoventral patterning, blastema cell proliferation and cartilage induction during axolotl tail regeneration. Development, 132(14), 3243–3253. 10.1242/dev.01906

Sehm, T., Sachse, C., Frenzel, C., & Echeverri, K. (2009). miR-196 is an essential early-stage regulator of tail regeneration, upstream of key spinal cord patterning events. Dev Biol, 334(2), 468–480. 10.1016/j.ydbio.2009.08.008

Showell, C., Binder, O., & Conlon, F. L. (2004). T-box genes in early embryogenesis. Dev Dyn, 229(1), 201–218. 10.1002/dvdy.10480

Slack, J. M., Beck, C. W., Gargioli, C., & Christen, B. (2004). Cellular and molecular mechanisms of regeneration in Xenopus. Philos Trans R Soc Lond B Biol Sci, 359(1445), 745–751. 10.1098/rstb.2004.1463

Sobkow, L., Epperlein, H. H., Herklotz, S., Straube, W. L., & Tanaka, E. M. (2006). A germline GFP transgenic axolotl and its use to track cell fate: dual origin of the fin mesenchyme during development and the fate of blood cells during regeneration. Dev Biol, 290(2), 386–397. 10.1016/j.ydbio.2005.11.037

Stewart, S., Tsun, Z. Y., & Izpisua Belmonte, J. C. (2009). A histone demethylase is necessary for regeneration in zebrafish. Proc Natl Acad Sci U S A, 106(47), 19889–19894. 10.1073/pnas.0904132106

Svetlov, P. (1934). Über die Regeneration während der Embryonalentwicklung: Wiederherstellung des Schwanzes und der Schwanzknospe bei Rana temporaria. Wilhelm Roux’Archiv für Entwicklungsmechanik der Organismen, 131(4), 672–701.

Taniguchi, Y., Kurth, T., Weiche, S., Reichelt, S., Tazaki, A., Perike, S., Kappert, V., & Epperlein, H. H. (2017). The posterior neural plate in axolotl gives rise to neural tube or turns anteriorly to form somites of the tail and posterior trunk. Dev Biol, 422(2), 155–170. 10.1016/j.ydbio.2016.12.023

Tucker, A. S., & Slack, J. M. W. (1995). The Xenopus laevis tail-forming region. Development, 121(1), 249–262. 10.1242/dev.121.1.249

Vincent, C. D., Rost, F., Masselink, W., Brusch, L., & Tanaka, E. M. (2015). Cellular dynamics underlying regeneration of appropriate segment number during axolotl tail regeneration. BMC Dev Biol, 15, 48. 10.1186/s12861-015-0098-1

Vogt, W. (1931). Über regeneratives und regulatives Wachstum: nach Defektversuchen an Schwanz und Schwanzknospe von Amphibienkeimen. Verlag von Gustav Fischer.

Wang, L., Song, L., Yi, C., Zhou, J., Yong, Z., Hu, Y., Pan, X., Qiao, N., Cai, H., Zhao, W., Zhang, R., Yang, L., Liu, L., Peng, G., Tanaka, E. M., Li, H., Liu, Y., & Fei, J. F. (2026). Divergent stem cell mechanisms govern the primary body axis and appendage regeneration in the axolotl. Sci Adv, 12(6), eadx5697. 10.1126/sciadv.adx5697

Wells, K. M., Kelley, K., Baumel, M., Vieira, W. A., & McCusker, C. D. (2021). Neural control of growth and size in the axolotl limb regenerate. Elife, 10. 10.7554/eLife.68584

Williams, M. C., Patel, J. H., Kakebeen, A. D., & Wills, A. E. (2021). Nutrient availability contributes to a graded refractory period for regeneration in Xenopus tropicalis. Dev Biol, 473, 59–70. 10.1016/j.ydbio.2021.01.005

Wilson, V., Manson, L., Skarnes, W. C., & Beddington, R. S. (1995). The T gene is necessary for normal mesodermal morphogenetic cell movements during gastrulation. Development, 121(3), 877–886. 10.1242/dev.121.3.877

Woodcock, M. R., Vaughn-Wolfe, J., Elias, A., Kump, D. K., Kendall, K. D., Timoshevskaya, N., Timoshevskiy, V., Perry, D. W., Smith, J. J., Spiewak, J. E., Parichy, D. M., & Voss, S. R. (2017). Identification of Mutant Genes and Introgressed Tiger Salamander DNA in the Laboratory Axolotl, Ambystoma mexicanum. Sci Rep, 7(1), 6. 10.1038/s41598-017-00059-1

Wymeersch, F. J., Skylaki, S., Huang, Y., Watson, J. A., Economou, C., Marek-Johnston, C., Tomlinson, S. R., & Wilson, V. (2019). Transcriptionally dynamic progenitor populations organised around a stable niche drive axial patterning. Development, 146(1). 10.1242/dev.168161

Wymeersch, F. J., Wilson, V., & Tsakiridis, A. (2021). Understanding axial progenitor biology in vivo and in vitro. Development, 148(4). 10.1242/dev.180612

